# Spatial and single-cell transcriptomics capture two distinct cell states in plant immunity

**DOI:** 10.1101/2025.06.03.657683

**Authors:** Yuzhao Hu, Raeann Schaefer, Michael Rendleman, Andrew Slattery, Annaliese Cramer, Abdullah Nahiyan, Lori Breitweiser, Mokshada Shah, Emma Kaehler, Chenglin Yao, Andrew Bowling, John Crow, Gregory May, Girma Tabor, Shawn Thatcher, Srinivasa Rao Uppalapati, Usha Muppirala, Stéphane Deschamps

## Abstract

Unlike animals, plants are sessile organisms that must adapt to localized and fluctuating environmental stimuli, including abiotic and biotic stresses. While animals use mobile immune cells to eliminate pathogens, plants rely on localized cells in contact with the pathogen to detect and mount immune responses. Although bulk RNA sequencing (RNA-seq) has enabled the assessment of plant responses to pathogen infection at the whole transcriptome level, the spatial coordination of plant immune responses remains elusive. In this study, we performed both spatial and single-nuclei transcriptomic experiments to capture the spatial pattern of soybean plant responses to Asian soybean rust infection caused by the pathogen *Phakopsora pachyrhizi*. Through the analysis of both spatial and single-nuclei transcriptomics data, we identified two distinct host cell states with specific spatial localization in response to pathogen infection: the infected regions with the presence of the pathogen and the surrounding regions bordering the infected regions. Importantly, the surrounding regions exhibited higher expression of defense response-related genes than the infected regions, despite having minimal presence of the pathogen, indicating a cell non-autonomous defense response in the surrounding regions. Additionally, gene co-expression network analysis with single-cell resolution identified a key immune response-related gene module activated in the stressed cells captured in our single-nuclei RNA-seq data. This study reveals the intricate spatial coordination of plant defense responses against pathogen infection and enhances our understanding of the importance of localized cell non-autonomous defense responses in plant-pathogen interactions.

## 1 Introduction

Plant pathogens represent a serious threat to crop production worldwide. Historically, the agricultural management of diseases has focused on multi-pronged approaches, including the use of pesticides, and adapted agronomic practices. More recent efforts have converged on the discovery of biotechnological solutions through the characterization of genes involved in resistance to specific pathogens. However, continuous adaptation of pathogens to their environment and the resulting emergence of resistance to treatments are well-known phenomena that require the ongoing discovery of new resistant genes to create durable and broad-spectrum resistance.

The mechanisms involved in a plant’s response to pathogen infection are generally well-known (Ngou et al., 2021). The first layer involves cell surface pattern recognition receptors (PRR) that recognize potential pathogens in the environment resulting in pattern-triggered immunity (PTI) (Dodds and Rathjen, 2010; Ngou et al., 2022). In turn, PTI initiates downstream molecular responses including the production of reactive oxygen species, calcium influx, and hormone production (Aerts et al., 2022). Pathogens, however, have evolved a series of small molecules, called effectors, which, when injected into host cells, interfere with PTI. To counteract this effect, another class of intracellular immune receptors, nucleotide-binding domain leucine-rich repeat-containing receptors (NLRs), detect the cytoplasmic effectors delivered by the pathogen and activate effector-triggered immunity (ETI) (Cui et al., 2015). Both PTI and ETI initiate massive transcriptional reprogramming. Even with significant advancements, the cellular mechanisms and pathways triggered by PTI and ETI are not completely understood, and their investigation is often challenged by the fact that pathogen interaction with plant hosts can be highly heterogeneous (Latijnhouwers et al., 2003; Fawke et al., 2015). The host response is generally influenced by the distribution of the pathogen in the plant and its developmental changes during the colonization of the tissue (O’Connell et al., 2012). Those variable interactions can lead to asynchronous cellular interactions and trigger gene regulatory functions that can be specific to a particular cell type or cell state. Therefore, to better understand the host transcriptional responses to variable stages of infection, it is important to account for the spatial attributes and cell type-specific features of plant defense mechanisms and interactions with a pathogen.

Several recent studies have explored the underlying mechanisms of plant immunity at the single-cell level. Tang et al. used single-cell transcriptomics to determine the cell type-specific response of *Arabidopsis thaliana* leaf protoplasts to fungal pathogen *Colletotrichum higginsianum* (Tang et al., 2023). They demonstrated that immune receptor gene expression was highly heterogeneous across distinct cell types and included an enrichment of intracellular immune receptors in vasculature cells. Zhu et al. similarly explored the response of *A. thaliana* leaf protoplasts to infection with virulent *Pseudomonas syringae* using single-cell transcriptomics (Zhu et al., 2023b). They discovered cluster-specific gene expression patterns indicative of distinct cell states ranging from immunity to susceptibility whose nature depended on their proximity to the bacterial pathogen and the timeline of bacterial infection. More recently, Nobori et al. combined single-cell transcriptomics, epigenomics and spatial transcriptomics in *A. thaliana* infected with virulent and avirulent *P. syringae* to identify novel cell states in the host, in response to infection (Nobori et al., 2025). Specifically, they demonstrated the existence of a new cell state, labeled as “PRIMER” (primary immune responder) cells, located in regions of infection involved in the host ETI response and classified them, based on pseudotime analysis, as early responders to the bacterial infection. Additionally, PRIMER cells were surrounded by yet another novel cell state in response to infection, labeled as “bystander” cells, whose spatial and molecular profiles suggested they were involved in cell-to-cell immune signaling (Nobori et al., 2025). Those findings are in alignment with a perspective offered by Jacob et al, where ETI does not inhibit pathogen growth inside the initial infection site but rather induces cells that are not in direct contact with the pathogen through a process similar to the hypersensitive localized acquired resistance (LAR) immune response (Jacob et al., 2023a).

Taken together, these studies demonstrate the ability of single-cell and spatial techniques to characterize rare and localized cell states involved in early ETI response mechanisms (Delannoy et al., 2023; Tang et al., 2023; Zhu et al., 2023a), which may include genes involved in triggering multimodal resistance pathways, such as transcription factors. Similarly, pathogens infecting plants trigger and counteract plant resistance responses, by deploying a variety of strategies aimed, for example, at facilitating cell penetration or directing nutrient release (Langenbach et al., 2016). Such manipulation generally requires the interaction of pathogen effectors to plant proteins that are typically involved in normal growth and maintenance aspects of the plant metabolism. Therefore, both the infection of a plant by a pathogen and its immune response requires the redirection, either activation or suppression, of specific plant gene expression.

*Phakopsora pachyrhizi* is an obligate biotrophic fungus known to be the causative agent of Asian Soybean Rust (ASR) (Chicowski et al., 2024). Left untreated, *P. pachyrhizi* can defoliate soybean fields within a few days and trigger yield losses of up to 90%. It is especially prevalent in South America, and especially in Brazil, where it can cause economic losses that can reach several billion USD. It is currently controlled through the combined use of fungicides with different modes of action. Genetic resistance is known but is not effective against all isolates of ASR (Godoy et al., 2016). A more detailed molecular understanding of the Soybean-ASR interactions at spatial and single-cell level may help us identify key effectors and susceptibility genes and develop novel durable disease management solutions.

In this study, we analyzed the spatial and temporal genic response of host cells in relation to their proximity to *P. pachyrhizi* and the extent of infection within different regions of the leaf tissue. Our findings indicate that host cells at the sites of pathogen infection exhibit progressively lower transcriptional defense response compared to host cells at the surrounding regions. On the other hand, cells in the surrounding non-infected regions exhibit stronger transcriptional defense response, in alignment with the “LAR” model of plant cell non-autonomous immune response.

## 2 Materials and Methods

### 2.1 Plant growth and pathogen inoculation

For 10x Genomics Visium experiments, 10-day old Williams 82 soybean plants were inoculated with *Phakopsora pachyrhizi* isolate FL-07 (90-110K spores/ml) and collected at 4 DPI and 7 DPI. For 10x Genomics Chromium experiments, 14-day old 93Y21 soybean seedlings were spray inoculated with *P. pachyrhizi* isolate GA-05 at a high spore concentration (∼100K spores/ml). Inoculated unifoliate leaves were collected at 1, 3, and 5 DPI.

### 2.2 Single-nuclei RNA-seq

#### 2.2.1 Tissue collection

For each single-nuclei RNA-seq (snRNA-seq) sample, about 40 mg of fresh ASR-infected soybean leaf tissue was flash frozen in liquid nitrogen and stored at −80°C until processed.

#### 2.2.2 Nuclei isolation

For nuclei isolation, a previously published protocol was used with modifications (Guillotin et al., 2023). In brief, frozen tissue was transferred onto a petri dish (Corning #3160-60) pre-cooled on ice with 300 μl lysis buffer consisting of 0.3 M sucrose, 1.25% Ficoll (Fisher Scientific #34-169-125GM), 2.5% Dextran 40 (Millipore Sigma #31389-25G), 15 mM Tris HCl (pH 8), 20 mM 2-(N-morpholino) ethanesulfonic acid (MES), 10 mM MgCl_2_, 60 mM KCl, 15 mM NaCl, 0.5 mM spermine, 0.5 mM spermidine, 0.1% Triton X-100, 5 mM dithiothreitol (DTT), 0.1 mM phenylmethylsulfonyl fluoride (PMSF), 0.1% (v/v) protease inhibitor cocktail (Millipore Sigma #P9599), 0.4% bovine serum albumin (BSA), and 0.4 U/μl RNase inhibitor (Fisher Scientific #NC0148451). From this step forward, the tissue and the subsequent extracted nuclei were kept on ice throughout the nuclei isolation process. Next, the tissue is chopped in the petri dish on ice for 8 to 10 minutes using a razor blade. The samples were then transferred to a Dounce tissue grinder (DWK Life Sciences #885302). 400 μl lysis buffer was then used to wash the petri dish and added to the Dounce tissue grinder. 10 movements up-down were performed using the A pestle for the Dounce tissue grinder and the samples were left incubating in the Dounce tissue grinder for 5 minutes before another 5 movements up-down were performed. Samples were then filtered through a 20 μm cell strainer (Sysmex # 04-004-2324) into a 1.5 ml tube. 500 μl lysis buffer was used to wash the Dounce tissue grinder, then filtered through the 20 μm cell strainer into the same 1.5 ml tube. Samples were then centrifuged at 500 g for 10 minutes at 4°C. Supernatant was removed from each tube and 60 μl final buffer (0.3 M sucrose, 15 mM Tris HCl (pH 8), 15 mM MES, 60 mM KCl, 15 mM NaCl, 0.5 mM spermine, 0.5 mM spermidine, 5 mM DTT, 0.1% (v/v) protease inhibitor cocktail, 1% BSA, 0.2 U/μl RNase inhibitor) was added to resuspend the nuclei pellet. Resuspended nuclei were filtered through a 10 μm cell strainer (PluriSelect #43-10010-50). Isolated nuclei were kept on ice until ready to proceed with 10x Chromium library preparation.

#### 2.2.3 10x Genomics Chromium library preparation

Isolated nuclei were counted using a hematocytometer and about ten thousand nuclei from each sample were loaded onto 10x Genomics Chromium Next GEM chips. 10x Genomics Chromium libraries were prepared following manufacturer’s instructions, using reagents from 10x Genomics Chromium Next GEM Single Cell 3 Reagent Kits v3.1 (10x Genomics #PN-1000121).

#### 2.2.4 Sequencing

Chromium Next GEM libraries were pooled and sequenced on Illumina NovaSeq 6000 using read length recommended by 10x Genomics (Read 1: 28 cycles; i7 index: 10 cycles; i5 index: 10 cycles; Read 2: 90 cycles) and the sequencing depth of 300 million clusters per sample.

### 2.3 Visium spatial transcriptomics

#### 2.3.1 Tissue collection

Leaves were dissected from the plants and dissected further into small rectangles, placed in embedding molds (Ted Pella #27183) filled with Sakura Optimal Cutting Temperature (OCT) embedding medium (Ted Pella #27050) and oriented in cross section or paradermal using a straight pin. Molds were flash frozen in liquid nitrogen-chilled isopentane and stored at −80°C until further processing.

#### 2.3.2 Cryo-sectioning and fluorescence imaging

Sample blocks were moved to Leica CM1950 Cryostat and chamber temperature was set to −13°C. Blocks were allowed to acclimate and 10 μm-thick sections were cut and observed on Leica DM1000 to check for fungal structures. To increase the integrity of cryosections, the plastering technique from Dr. Stephen Peters was modified, using 1:1 diluted OCT in 10 mM phosphate-buffered saline (PBS) (Peters, 2003a; b). Sections were stained overnight at 4°C using wheat germ agglutinin Alexa Fluor 488 conjugate (Invitrogen #W11261) in PBS at 1:2000, mounted in 80% glycerol, and imaged using a 20x air objective on a Leica Upright DM5000 in fluorescence mode, using filter sets A4, Cy5, and GFP. After fungal structures were confirmed with staining, 10 μm-thick sections were collected onto 10x Genomics Visium slides: sections were individually cut, placed in a capture area on the slide, and melted to the slide using the tip of a finger applied to the side opposite the tissue while still in the cryostat chamber. Section and slide were immediately refrozen in cryostat chamber by placing the slide on the stage, and sectioning continued in this manner to complete all capture areas on the slide. Slides were stored at −80°C until further processing. After sectioning for the Visium slide, one additional post-processing section was generated and stained and imaged using the methods mentioned above to check for fungal structures again.

#### 2.3.3 Section fixation, staining and imaging

The 10x Genomics Visium slide with sections was removed from the −80°C and placed on a 37°C slide warmer for 1 minute. The slide was then placed on a plastic tray, and the tray was put on ice for the following steps. Farmer’s fixative (3-parts 100% Ethanol: 1-part Glacial Acetic Acid) was chilled to −20°C and placed on ice. Approximately 1 ml of fixative was pipetted onto the slide and left to stand for 2 minutes. The fixative was removed by tipping the slide. Approximately 1 ml of 100% ethanol, chilled to −20°C then placed on ice, was pipetted onto the slide and left to stand for 2 minutes. The ethanol was tipped off and the slide was allowed to dry for 2 minutes. Toluidine blue 1% in 1% Sodium Borate solution (Poly-Sciences #S2410-320Z) was diluted 1:10 with water, 1 ml pipetted on to the slide, and left to stand for 2 minutes. The stain was tipped off, 1 ml distilled water was pipetted to the slide and left to stand for 2 minutes. After the water was removed, the slide was allowed to dry for 5 minutes before imaging. The slide was imaged with transmitted light on a Leica Thunder Imager (Leica Microsystems) using a 10x/0.30NA air objective and a K5 monochrome camera. The slide was then placed on ice until the next steps.

#### 2.3.4 10x Genomics Visium tissue optimization

Tissue optimization was adapted from manufacturer’s instructions (CG000238 Rev F). 10 μm-thick sections were placed on Visium Tissue Optimization slides (10x Genomics), processed and imaged as indicated in section 2.3.3, and then were tested for various permeabilization times (5 minutes to 15 minutes). Reverse-transcription and tissue removal were performed using reagents from the Visium Spatial Tissue Optimization Reagent kits (10x Genomics #PN-1000193). After processing, the slide was imaged on a Leica Thunder Imager (Leica Microsystems) using a 10x/0.30NA air objective and a K5 monochrome camera. The LED3 system and the filter TXR (Excitation: BP 560/40, Dichroic: LP 585, Emission: BP 630/75) were used. The exposure time was set using the RNA control well as a positive control. After fluorescence imaging of the sections, 10 min permeabilization times were selected based on maximum fluorescence signal with the lowest signal diffusion. This permeabilization time was used for subsequent handling of Visium Gene Expression slides.

#### 2.3.5 10x Genomics Visium gene expression library preparation

Fixed and stained 10 μm thick sections on Visium Gene Expression slides (10x Genomics) were processed according to manufacturer’s instructions, using reagents from the Visium Spatial Gene Expression Reagent Kit (10x Genomics) and the Library Construction Kit (10x Genomics), for cDNA amplification and library construction, respectively. Briefly, for each section, 10-minute permeabilization at 37°C was followed by 45-minute reverse transcription at 53°C. After second strand synthesis and denaturation, cDNA fragments were recovered from individual sections and amplified for 17 cycles, as determined by the Cq value from the quantitative polymerase chain reaction (qPCR) optimization steps performed on the recovered cDNA fragments. After enzymatic fragmentation, end repair, adaptor ligation and size-selection, the resulting cDNA inserts were amplified using individual barcoded primers from the Dual Index TT Set A kit (10x Genomics), then pooled prior to sequencing.

#### 2.3.6 Sequencing

Sequencing was performed on Illumina NovaSeq 6000 and NovaSeq X systems according to 10x Genomics’ recommendations for read length (Read 1: 28 cycles; i7 index: 10 cycles; i5 index: 10 cycles; Read 2: 90 cycles) and depth (50,000 clusters per spot). Spatial localization and characterization of unique molecular events were performed using individual spatial barcode and unique molecular identifier (UMI) sequences derived from Read 1 sequencing data, while Read 2 sequencing data were used to determine the molecular identity (e.g., gene annotation) of the original cDNA fragment and transcript the sequence was derived from.

### 2.4 Processing and analysis of Visium spatial and Chromium snRNA-seq data

#### 2.4.1 Read mapping

The 10x Genomics Space Ranger software (v2.0.1) and 10x Genomics Cell Ranger software (v8.0.1) were used to process raw reads into count values for each barcoded Visium spot and barcoded chromium nuclei respectively (Zheng et al., 2017). For both experiments, reads were simultaneously aligned to the *Phakopsora pachyrhizi* genome (Phapa1, GCA_025201825.1) (Gupta et al., 2023) and *Glycine max* Williams 82 (Wm82v4) (Valliyodan et al., 2019). Data analysis for both the single-nuclei and spatial data was performed using the Seurat 5.1.0 package in R 4.3.2 (Satija et al., 2015).

#### 2.4.2 Sample filtering

Fiducial frames around the capture area on each Visium slide were aligned manually, and tissue-covered spots were selected using 10x Genomics Loupe Browser software (v8.1.2). After this, 33,994 genes (28,789 from soybean, 5,196 from ASR) across 5,085 spots in the twelve samples were used for further analysis.

High quality nuclei with *300 < UMI < 10,000; 250 < genes < 6000*; and less than 10% mitochondrial or chloroplastic reads were selected for further analysis. Doublets were identified and removed using DoubletFinder (v2.0.4) (McGinnis et al., 2019). After filtering, 44,478 genes (41,028 from soybean 3,450 from ASR) across 48,358 nuclei in the six samples were used for further analysis.

#### 2.4.3 Normalization and clustering

Spatial and single-nuclei samples were individually processed in the same manner yielding two distinct Seurat objects. Seurat *SCTransform* was used to normalize each sample before canonical correlation analysis (cca) integration of all capture areas or all single-nuclei samples (Hafemeister and Satija, 2019). Using the integrated data object, all spots or cells were clustered using 30 principal components (PCs) using the Louvain algorithm with a 0.5 resolution or 0.25 resolution for spatial and single-nuclei objects respectively. Uniform manifold approximation and projection (UMAP) dimensional reductions were also calculated with 30 PCs.

The stressed cell type cluster was subset from the full single-nuclei object where Louvain clustering was run with a resolution of 0.25. Following this, marker genes for the resulting subclusters were identified as below and compared to cell clusters identified in the full dataset. Preserving the original UMAP confirmed subcluster cell type annotation.

#### 2.4.4 Gene level analyses

Cluster marker genes were identified using Seurat function *FindAllMarkers* with significance defined by a Wilcoxon Rank Sum test and adjusted P < 0.05. Differentially expressed genes (DEGs) between the healthy and infected clusters were identified using Seurat function *FindMarkers* with significance defined by a Wilcoxon Rank Sum test and adjusted P < 0.05. Gene Ontology (GO) enrichment was determined with fgsea (v1.28.0) (Korotkevich et al., 2021) ranked by all *avg_log2FC* values calculated by the above functions. Seurat function *AddModuleScore* was used to summarize expression of 15 ASR induced genes originally identified by Cabre et al. (Cabre et al., 2021) and Morales et al (Morales et al., 2013).

#### 2.4.5 Normalized pseudo-bulk analysis

Raw counts were summed per gene for the spatial and single-nuclei datasets using the Seurat function *AggregateExpression*. The summed raw counts were supplied to DESeq2 (v1.42.1) for default normalization (Love et al., 2014).

#### 2.4.6 Visualizations

Several R packages were used to create data visualizations. EulerR (v7.0.2) and SuperExactTest (v1.1.0) were used to create Venn diagrams and the associated statistics (Wang et al., 2015; Larsson and Gustafsson, 2018). UpsetR (v1.4.0) was used to create Upset plots (Conway et al., 2017). Pheatmap (v1.0.12) was used to create all heatmaps (Kolde, 2018).

### 2.5 High dimensional weighted gene co-expression network analysis

High dimensional weighted gene co-expression network analysis (hdWGCNA) was performed using the hdWGCNA 0.3.03 package in R 4.3.3. Before network construction, genes were filtered to soybean genes expressed in at least 1% of cells (24,705 genes), and cells were filtered to those labeled as healthy or stressed mesophyll cells. The *ConstructNetwork* function was used to build signed co-expression networks with *minModuleSize* set to 50, *detectCutHeight* set to 0.995, and *mergeCutHeight* set to 0.2. To select the soft power threshold, a parameter sweep was performed with the *TestSoftPowers* function as described by Morabito et al (Morabito et al., 2023).

To summarize gene expression across the genes in each co-expression module, module eigengenes (MEs) were calculated. MEs are defined as the first principal component of the module’s gene expression matrix, and module hub genes are selected by correlation with the module’s eigengene. A gene-centric UMAP reduction was performed on the topological overlap of each gene with the top 10 hub genes from each module. For statistical comparison of module eigengene values, the unpaired two-sided Wilcoxon Rank Sum test was employed.

## 3 Results

### 3.1 Spatial and single nuclei transcriptomics simultaneously capture host and pathogen transcriptome

To characterize the spatial and temporal changes in the transcriptome during plant-pathogen interaction, we infected soybean unifoliate leaves with ASR spores (Figure 1A). We generated paradermal (PD) sections of soybean leaves at 4 days post-infection (DPI) and cross-sections (CS) of soybean leaves at both 4 DPI and 7 DPI (Figure 1A). For each sample type, four serial sections were used for 10x Genomics Visium experiments (Figure S1A). Once tissue sections were placed onto Visium slides, the RNAs were permeabilized onto the slides, which contain spots with primers that have spatial barcodes, UMIs, and poly(dT) primers to capture transcripts with poly(A) tails (Figure 1A) (Marx, 2021). The spatial barcodes allow tracing of sequencing reads back to specific spots on the Visium slide, which can then be overlaid and compared to brightfield images of the sections on the slides to obtain spatial information on gene expression (Marx, 2021). The poly(dT) primers enable the capture of poly(A) tailed RNA molecules (Verma, 1978), which are prevalent in both soybean and ASR nuclei (Fasken and Corbett, 2009), allowing the simultaneous capture of soybean and ASR transcriptomes in the spatial transcriptomics experiment.

**Figure 1.**
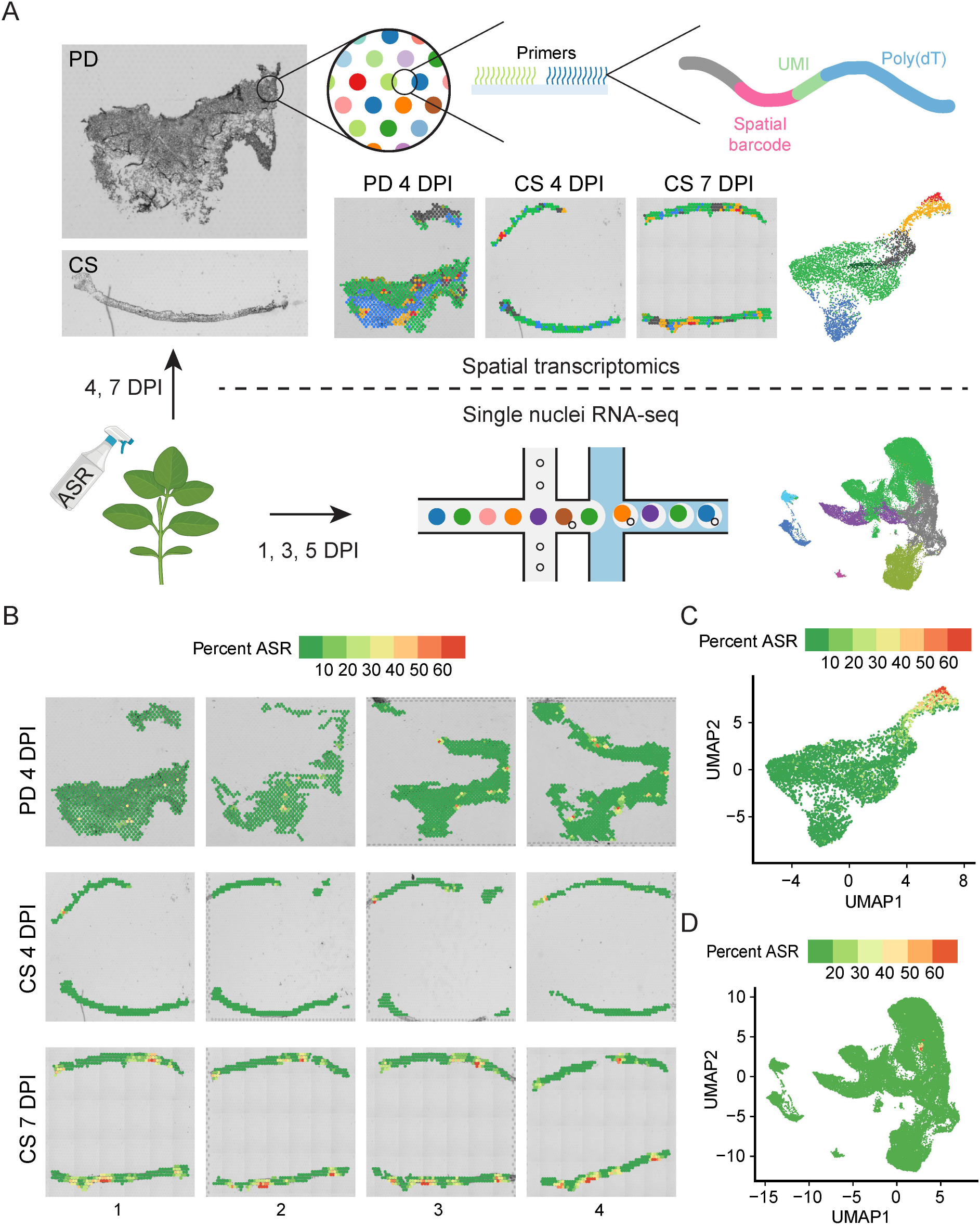
Spatial transcriptomics and snRNA-seq simultaneously capture both soybean and ASR transcriptome. **A)** Schematic of experiments performed for various samples of ASR-infected soybean leaf. Leaves infected for 4 days were paradermal (PD) and cross sectioned (CS), leaves infected for 7 days were cross sectioned for Visium spatial analysis. Leaves infected for 1, 3, and 5 days were subjected to snRNA-seq. **B)** Percent ASR reads per spot mapped spatially reveals localized regions of ASR infection. **C)** Spatial UMAP of percent ASR reads per spot. **D)** snRNA-seq UMAP of percent ASR reads per nuclei.

Total soybean genes captured from the 12 Visium sections ranged from nineteen thousand to thirty-two thousand and ASR genes ranged from one thousand to six thousand (Figure S1B), indicating sufficient coverage of the whole transcriptome of both soybean and ASR. Principal component analysis of all 12 Visium sections revealed that the 4 DPI and 7 DPI samples formed distinct clusters (Figure S1C), suggesting that the expression of ASR and soybean genes changes over the time course of infection. Additionally, we analyzed the percentage of ASR reads per spot and found that the ASR read percentage is significantly higher in 7 DPI cross-sections compared to either 4 DPI paradermal sections or 4 DPI cross-sections (Figure S1D), indicating the expansion of ASR infection in soybean leaves as the disease progresses. Spatial mapping of the percentage of ASR reads per spot shows spots with high ASR presence in each section (Figure 1B), highlighting the spatial heterogeneity of ASR infection in soybean leaves. To analyze the relationship between different spots across all sections, we combined all spots from the 12 sections and generated a UMAP (Figure 1C and S1E). After combining data from the different sections, the UMAP showed that spots from different sections were well integrated (Figure S1E). Furthermore, we examined the percentage of ASR reads per spot and found that spots with a high percentage of ASR reads localized closely together on the UMAP (Figure 1C), indicating that ASR-infected regions have similar gene expression profiles. Together, these results demonstrate that spatial transcriptomics can simultaneously capture both soybean and ASR transcriptomes and identify specific spatial regions of active ASR infection.

Although the spatial transcriptomic experiments capture gene expression along with *in situ* spatial localization information, the spots on the Visium slides have a resolution of 55 µm, which is not true single-cell resolution (Figure 1A). Therefore, we complemented the spatial transcriptomics study with snRNA-seq using the 10x Genomics Chromium platform (Figure 1A). We isolated nuclei from ASR-infected soybean leaves at 1, 3, and 5 DPI, with two biological replicates for each time point, and subjected the nuclei to snRNA-seq (Figure 1A). Across the six snRNA-seq samples, the number of soybean cells captured ranged from five thousand to eleven thousand, and the total number of soybean genes detected ranged from forty-two thousand to forty-six thousand (Figure S1F), suggesting that the snRNA-seq data captured most of the whole transcriptome of soybean cells. Further PCA analysis of all six snRNA-seq samples showed that all the biological replicates clustered together (Figure S1G), indicating the reproducibility of the snRNA-seq. Similar to the Visium platform, the Chromium platform also uses poly(dT) primers for capturing transcripts, allowing the simultaneous capture of soybean and ASR transcripts, as indicated by the capture of a total of three thousand to five thousand ASR genes in the six snRNA-seq samples (Figure S1F). Accordingly, the percentage of ASR reads per nucleus significantly increased from 1 DPI to 3 DPI and from 3 DPI to 5 DPI in the snRNA-seq samples (Figure S1H), confirming the progression of ASR infection over time. To explore the relationship between the gene expression of all nuclei captured in the six snRNA-seq samples, we generated a UMAP of all nuclei (Figure 1D and S1I). After combining the data, nuclei from different samples were evenly distributed across the UMAP (Figure S1I), indicating good integration and allowing for the examination of gene expression changes between different samples in specific nuclei clusters. In the snRNA-seq dataset, a small cluster of ASR nuclei showed a high percentage of ASR reads per nucleus (Figure 1D), again indicating the successful capture of both soybean and ASR transcriptomes in the snRNA-seq experiment.

Altogether, these results demonstrate that both spatial transcriptomics and snRNA-seq can simultaneously capture the transcriptomes of both the host soybean and the pathogen ASR. The spatial transcriptomics data allows for the visualization of the ASR infection center, highlighting the heterogeneity of pathogen infection in plants. Meanwhile, the snRNA-seq data complements the spatial transcriptomics data with single-nuclei resolution, enabling cell-specific analysis of gene expression changes upon pathogen infection.

### 3.2 Single-nuclei RNA-seq captures a stressed cell cluster

Pathogen infection in plants often results in heterogeneous responses among different host cells in the infected organ, as characterized in previous single-cell transcriptomic studies focusing on plant-pathogen interactions (Cao et al., 2023; Delannoy et al., 2023; Tang et al., 2023; Nobori et al., 2025; Yan et al., 2025). Single-cell/nuclei RNA-seq provides unique opportunities to identify different cell types and even different cell states in the host plant in response to pathogen infection (Zhu et al., 2023b). To elucidate the differential transcriptomic responses among various host cell types/states during pathogen infection, we performed clustering of all nuclei assayed by snRNA-seq and identified 12 distinct clusters, labeled from cluster 0 to 11 (Figure S2A). It is known in the literature that soybean homologs of cell type marker genes from other plant species often exhibit conserved expression in the same soybean cell types (Cervantes-Pérez et al., 2024). Therefore, to annotate the nuclei clusters in our snRNA-seq data, we searched the literature and the Single Cell Plant Database (scPlantDB) for known leaf cell type marker genes in soybean and other plant species and checked the expression of these marker genes or their soybean homologs in our clusters (He et al., 2024). Based on these known leaf cell type marker genes, we annotated a majority of the clusters in our snRNA-seq data: clusters 0, 2, 6, and 9 as mesophyll cells; cluster 1 as pavement cells; cluster 4 as bundle sheath cells; cluster 7 as phloem parenchyma cells; cluster 8 as phloem companion cells; and cluster 10 as guard cells (Table 1, Figure 2A).

**Figure 2.**
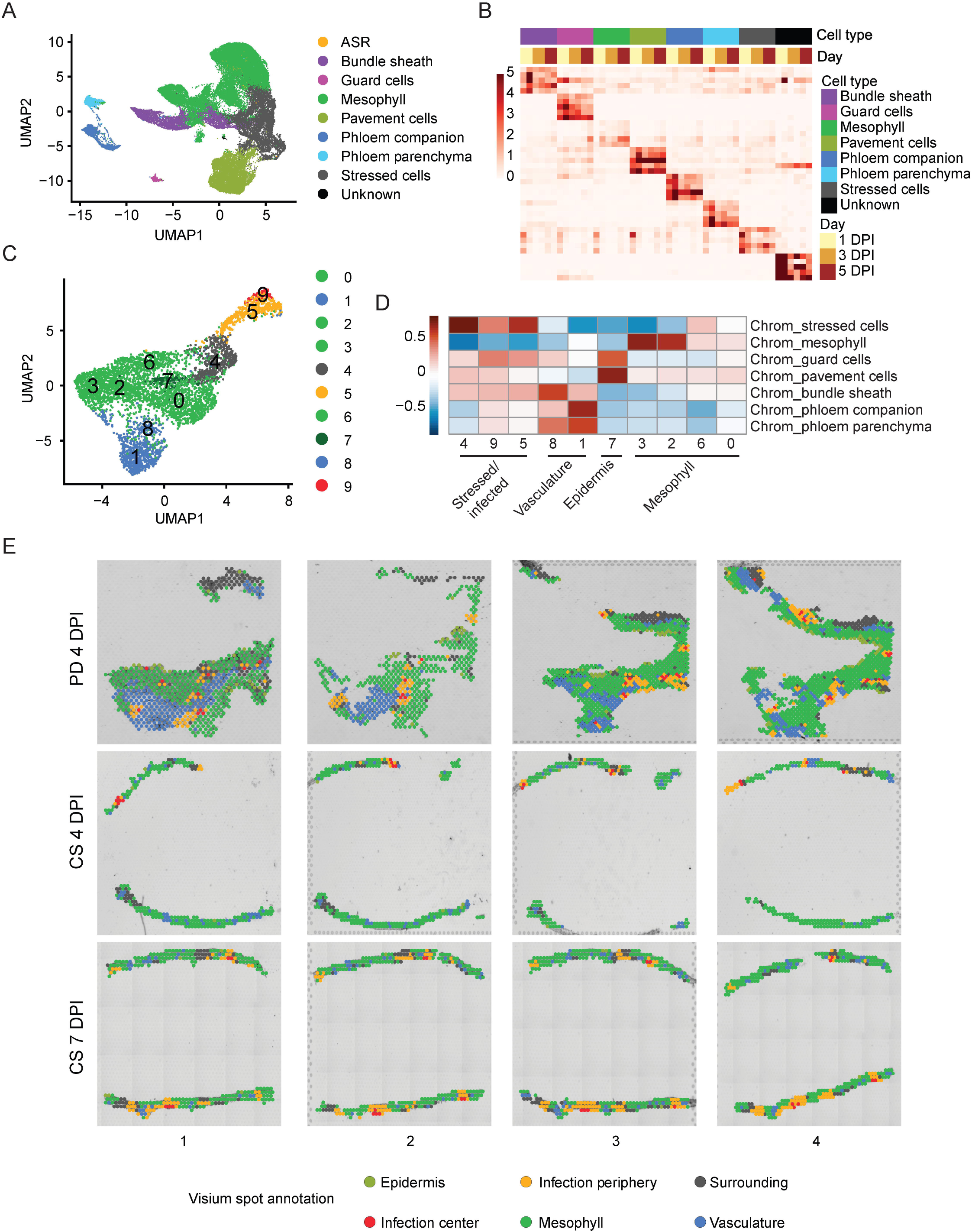
Annotation of snRNA-seq and spatial clusters. **A)** UMAP of annotated snRNA-seq clusters. **B)** heatmap of normalized pseudo-bulked expression of top marker genes for each annotated cell type within snRNA-seq. **C)** UMAP of Spatial clusters. **D)** Correlation heatmap of marker genes in annotated snRNA-seq cell types and spatial clusters. **E)** Annotation of all spots for spatial transcriptomics data.

**Table 1.** Summary of known leaf cell type marker genes used to annotate snRNA-seq clusters.

| Cell type | *Arabidopsis* marker gene | Soybean marker gene | snRNA-seq cluster | Reference |
| --- | --- | --- | --- | --- |
| Bundle sheath | *AT3G51895* | *Glyma.10G028900* | 4 | (Kim et al., 2021) |
| Guard cell | *AT3G24140* | *Glyma.02G129100* | 10 | (Kim et al., 2021) |
| Pavement cell | *AT1G27950* | *Glyma.18G005800* | 1 | (Kim et al., 2021) |
| Phloem<br>parenchyma cell | *AT3G56620* | *Glyma.03G122200* | 7 | (Liu et al., 2022) |
| Phloem<br>companion cell |  | *Glyma.07G222500* | 8 | (Cervantes-Pérez et al., 2024) |
| Mesophyll cell | *AT3G01500* | *Glyma.05g007100* | 0, 2, 6, 9 | (Procko et al., 2022) |

However, clusters 3 and 5 from the snRNA-seq dataset did not show enriched expression of any of the previously mentioned known leaf cell type marker genes and therefore could not be annotated using this approach. Since pathogen infection can induce drastic gene expression changes in host cells and obscure their cell type identities (Tang et al., 2023), we derived cluster-specific marker genes from clusters 3 and 5 and performed GO term analysis using these cluster marker genes (Figure S2C, Table S1). Interestingly, marker genes from both clusters 3 and 5 showed enrichment in GO terms related to plant immune response (Figure S2C, Table S1). This result suggests that clusters 3 and 5 are immune-responsive or stressed cells, likely caused by ASR infection. Therefore, we annotated clusters 3 and 5 together as stressed cells (Figure 2A). To verify if clusters 3 and 5 are indeed immune-responsive cells in response to ASR infection, we curated a list of ASR-induced genes from previous bulk transcriptome studies performed using ASR-infected soybean leaves (Table S2) (Morales et al., 2013; Cabre et al., 2021). To check the expression of these ASR-induced genes, we treated these genes as an ASR-induced gene module and calculated the expression of this module across all nuclei (Figure S2D, Table S2). Interestingly, the ASR-induced gene module expression is mostly enriched in clusters 3 and 5 (Figure S2D), suggesting that these stressed cells exhibit transcriptional changes in response to ASR infection. Apart from the clusters we have annotated, cluster 11 had the fewest number of cells and did not show specific cell type marker gene expression, so we designated cluster 11 as an unknown cluster and excluded it from the rest of the analysis (Figure 2A and S2A). Moreover, in our snRNA-seq data, there are ASR cells assigned based on the mapping of the majority of reads within the cell barcode to the ASR genome, so we designated those cells as ASR cells instead of one of the annotated soybean cell types (Figure 2A). Since we detected fewer than 100 ASR cells (Figure S1F and 2A), we also excluded the ASR cells from the rest of our snRNA-seq analysis to focus on host responses.

After annotating most of the nuclei clusters, we also examined the soybean cell type composition at different time points: 1 DPI, 3 DPI, and 5 DPI (Figure S2E). Furthermore, we identified cell type-specific marker genes for the annotated cell types and demonstrated that the expression of these marker genes is highly enriched in the corresponding cell types (Figure 2B). This will be a valuable resource for annotating cell clusters in future soybean single-cell transcriptomic studies. Altogether, these results suggest that our snRNA-seq data captures not only the major cell types in soybean leaves but also a specific cluster of stressed cells that are responsive to ASR infection.

### 3.3 Spatial transcriptomics identify sites of pathogen infection

To gain more insight into the spatial organization of the plant immune response to pathogens, we next sought to cluster the different spatial spots from all 12 sections and to annotate the spot clusters. We performed clustering analysis for all spots and identified 10 different spatial clusters, labeled from 0 to 9 (Figure 2C). Since we had already annotated the snRNA-seq clusters and derived cell type-specific marker genes for soybean leaves infected with ASR (Figure 2A-B), we correlated marker genes of the spatial clusters with the cell type-specific marker genes to facilitate the annotation of the spatial clusters (Figure 2D). Interestingly, spatial clusters 3, 2, 6, and 0 clustered strongly with mesophyll marker genes derived from the snRNA-seq performed in this study (Figure 2D), suggesting that these four spatial clusters can be annotated as the mesophyll region (Figure 2D-E). Moreover, spatial cluster 7 had the highest correlation with marker genes from guard cells and pavement cells (Figure 2D), indicating that spatial cluster 7 corresponds to the epidermis region (Figure 2D-E). Similarly, spatial clusters 8 and 1 strongly correlated with the marker genes identified for vasculature-related cells, including phloem companion cells, phloem parenchyma cells, and bundle sheath cells (Figure 2D), indicating that these spatial clusters correspond to the vasculature region (Figure 2D-E). Finally, spatial clusters 4, 9, and 5 had high correlation with marker genes from the stressed cells identified in the snRNA-seq experiment in this study (Figure 2D), suggesting that these spatial clusters are the stressed or infected regions (Figure 2D).

By overlaying the spatial clusters with the brightfield image for the sections, we visually confirmed the spatial clusters that correspond to the vasculature and epidermis regions (Figure S2F). The vasculature-related spatial clusters, clusters 1 and 8, are localized to the vasculature regions shown in both the paradermal sections and the cross sections (Figure S2F). Similarly, the epidermis-related spatial cluster, cluster 7, clearly localized to the epidermis regions, which is especially evident in the paradermal sections (Figure S2F). These results further validate the spatial cluster annotations. When we overlaid all annotated spatial clusters on the brightfield image of the sections, it further showed that the stressed/infected clusters, clusters 9, 5, and 4, had a specific spatial organization pattern: cluster 9 forming the infection center, cluster 5 forming the infection periphery that surrounds the infection center, and cluster 4 forming the surrounding region that further borders the infection center and infection periphery regions (Figure 2E). After annotating the spatial clusters using the transcriptomics data, we also sought to independently verify this annotation by staining the adjacent serial section of the same tissue block after generating the sections for spatial transcriptomics experiment. For the tissue block used for the 7 DPI cross-sections, we stained the post-processing section with wheat germ agglutinin Alexa Fluor 488 conjugate and imaged the section using a fluorescence microscope (Figure S2G). We found that the ASR cells indicated by green fluorescence overlapped with the infection center regions identified in the spatial transcriptomics data (Figure S2G). This result suggests that the annotated infection center regions using the snRNA-seq and spatial transcriptomics data agree with the visualized pathogen infection sites. Altogether, the spatial transcriptomics experiment performed in this study revealed the spatial organization of the host response to pathogen infection and further underscored the heterogeneity in plant-pathogen interactions.

### 3.4 Single-nuclei RNA-seq and spatial transcriptomics reveal key pathways in host response to pathogen infection

It is known that pathogen infection can induce defense-related gene expression changes in different host cell types and obscure cell identity in single-cell RNA-seq experiments, but sub-clustering can disentangle and further annotate obscured cell clusters (Tang et al., 2023). We have already annotated a cluster of stressed cells that showed immune response-related activity (Figure 2A and 3A). To unravel cell type identity, we performed sub-clustering for the stressed cells (Figure 3B), considering their relative relationship with the rest of the cells in the UMAP, and generated seven sub-clusters, labeled from sub-cluster 0 to 6 (Figure 3B). To further annotate these sub-clusters, we correlated their marker gene expression with the marker genes of the annotated cell types in the snRNA-seq data (Figure 3C). Sub-clusters 0, 1, and 2 correlated strongly with mesophyll cells (Figure 3C), suggesting that these sub-clusters correspond to stressed mesophyll cells (Figure 3C-D). Similarly, sub-clusters 4 and 3 had high correlations with bundle sheath cells (Figure 3C), prompting us to annotate these sub-clusters as stressed bundle sheath cells (Figure 3C-D). Finally, sub-clusters 5 and 6 had strong correlations with pavement cells (Figure 3C), suggesting that these sub-clusters are stressed pavement cells (Figure 3C-D).

**Figure 3.**
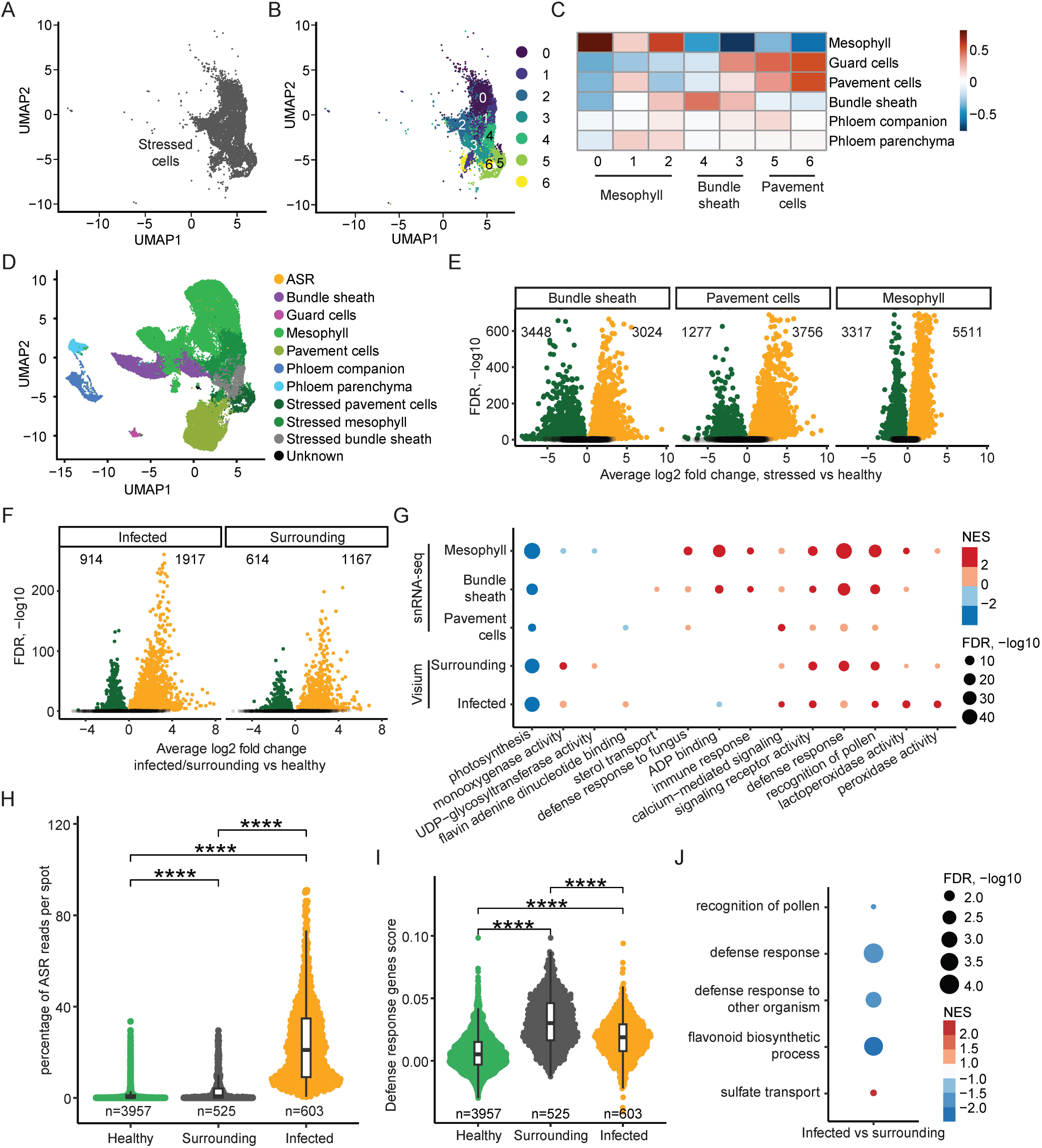
ASR Infection induces two spatially distinct response programs. **A)** UMAP of stressed cell cluster in snRNA-seq dataset. **B)** UMAP of subclustered stressed cells in snRNA-seq. **C)** Correlation heatmap of marker genes in annotated snRNA-seq clusters and sub-clusters of stressed cells. **D)** UMAP of snRNA-seq annotated clusters including annotated subclusters of stressed cells. **E)** Volcano plots of significantly differentially expressed soybean genes (adjusted *p* value < 0.05) in each stressed cell type compared to its corresponding healthy cluster. **F)** Volcano plots of significantly differentially expressed soybean genes (adjusted *p* value < 0.05) in each **spatial** region compared to the remaining healthy spots. **G)** Dot plot of enriched GO terms associated with differentially expressed genes from each comparison using both snRNA-seq and spatial transcriptomics data. Size of dots corresponds to −log10 of adjusted *p* value, colored by NES. **H)** Percent ASR reads per spot in each annotated **spatial** region. * indicates *p* < 0.05, ** indicates *p* > 0.01, *** indicates *p* < 0.001, **** indicates *p* < 0.0001. **I)** Defense response GO term-associated gene expression described as a single score per spot in each **spatial** region. * indicates *p* < 0.05, ** indicates *p* > 0.01, *** indicates *p* < 0.001, **** indicates *p* < 0.0001. **J)** Dot plot of enriched GO terms associated with genes differentially expressed between the infected and surrounding spatial regions. Size of dots corresponds to −log10 of adjusted *p* value, colored by NES.

After annotating the sub-clusters for the stressed cells, the UMAP of all annotated cells further validated the sub-cluster annotations, with the stressed mesophyll cells localizing close to the mesophyll cells, and similarly for the stressed bundle sheath cells and the stressed pavement cells (Figure 3D). This annotation of the sub-clusters of the stressed cells allowed us to perform DEG analysis by comparing the stressed mesophyll cells to the healthy mesophyll cells, the stressed bundle sheath cells to the healthy bundle sheath cells, and the stressed pavement cells to the healthy pavement cells (Figure 3E). After identifying the DEGs for mesophyll, bundle sheath, and pavement cells, we also performed GO term enrichment analysis for these three cell types and compared the GO terms (Figure S3A). By overlapping the significantly enriched GO terms for all three cell types, we found significant overlap between the mesophyll, bundle sheath, and pavement cells (Figure S3A), suggesting that different host cell types can activate similar defense response pathways in reaction to pathogen infection. Moreover, we also checked the expression of ASR-induced genes from the literature in stressed mesophyll cells versus healthy mesophyll cells, stressed pavement cells versus healthy pavement cells, and stressed bundle sheath cells versus healthy bundle sheath cells (Figure S3B, Table S2) (Morales et al., 2013; Cabre et al., 2021). We found that the ASR-induced genes had increased expression in the stressed cells from all three cell types (Figure S3B), indicating that pathogen infection could induce similar gene expression changes in different host cell types. different cell types.

Similar to the snRNA-seq, we also aimed to decipher the gene expression changes upon pathogen infection in the spatial data. As elucidated earlier in this study, there are three spatial clusters (clusters 4, 5 and 9) that correlated strongly with the stressed cells identified in the snRNA-seq (Figure 2C-D), suggesting that these three spatial clusters are related to the host response to the pathogen. Among these spatial clusters, spatial clusters 9 and 5 are closely localized on the spatial UMAP (Figure 2C), with spatial cluster 4 being further away from spatial clusters 9 and 5 (Figure 2C). Importantly, the percentage of ASR reads per spot are much higher in spatial cluster 9 and 5, compared to spatial cluster 4 (Figure 1C and 2C). Therefore, we combined the annotation of spatial cluster 9 (infection center) and spatial cluster 5 (infection periphery) together and annotated them as the infected region (Figure 2C, 2E and 3F), while spatial cluster 4 was annotated as the surrounding region (Figure 2C, 2E and 3F). Spatially, the surrounding regions are bordering the infected regions (Figure 2E). Next, we divided all spots into three categories: the infected region, the surrounding region, and the healthy region, and performed DEG analysis by comparing the infected region to the healthy region, as well as comparing the surrounding region to the healthy region (Figure 3F). We then performed GO term analysis using the DEGs identified in the infected and surrounding regions and overlapped the significantly enriched GO terms (Figure S3C). We found significant overlap in the GO terms for both the increased and decreased genes between the infected regions and surrounding regions (Figure S3C), suggesting that similar transcriptomic pathways are perturbed in the infected and surrounding regions compared to the healthy regions.

Furthermore, we examined the GO terms derived from DEGs from both the snRNA-seq and the spatial transcriptomics data and found many overlapping GO terms between these two datasets (Figure 3G and S3D, Table S1), suggesting that spatial and single-cell transcriptomics capture similar transcriptional changes in the host response to pathogen infection. Interestingly, the GO term “photosynthesis” was shown in the downregulated genes in all five comparisons: stressed mesophyll versus healthy mesophyll, stressed bundle sheath versus healthy bundle sheath, stressed pavement cells versus healthy pavement cells, surrounding region versus healthy region, and infected region versus healthy region (Figure 3G, Table S1). This aligns with previous studies reporting a decrease in photosynthesis activity in pathogen-infected host leaves (Bilgin et al., 2010). Similarly, immune response-related GO terms, such as “calcium-mediated signaling,” “signaling receptor activity,” and “defense response,” were enriched in the upregulated genes in all five comparisons from both snRNA-seq and spatial transcriptomics data (Figure 3G, Table S1), validating our findings that the stressed cells in the snRNA-seq and the surrounding and infected regions in the spatial transcriptomics data are all responsive to ASR infection. Altogether, these results show that both snRNA-seq and spatial transcriptomics reveal the key immune response pathways activated in host cells in response to pathogen infection.

### 3.5 Infection surrounding regions have minimal pathogen presence yet strong defense response

Upon examination of the GO term dot plot of DEGs in the spatial data, we noticed that the normalized enrichment score (NES) value for the GO term “defense response” is higher in the surrounding region compared to the infected region (Figure 3G, Table S1). However, as discussed earlier in this study, the infected region had a higher ASR read content compared to the surrounding region (Figure 1C, 2C and 2E). This result suggests that the surrounding region identified in the spatial transcriptomics data has less ASR presence but potentially a higher defense response compared to the infected regions. This is reminiscent of localized acquired resistance (Ross, 1961; Jacob et al., 2023a). It has been observed in the literature that in *Arabidopsis* leaves infected with the bacteria *Pseudomonas syringae pv. tomato DC3000 AvrRps4*, the defense-related gene *PR1* is highly expressed in the surrounding regions of the pathogen infection site but is rather repressed in the infected leaf region with pathogen presence (Jacob et al., 2023a; Jacob et al., 2023b). In the same study, the authors proposed that in the pathogen-infected regions, the immune response of the host cells is suppressed by pathogen-derived effectors, while in the surrounding regions, the host cells activate a cell non-autonomous immune response, with high defense response activity and yet low presence of the pathogen (Jacob et al., 2023a; Jacob et al., 2023b). Therefore, we sought to examine whether the surrounding and infected regions identified in our spatial transcriptomic study exhibit similar characteristics.

We first compared the percentage of ASR reads among the healthy, surrounding, and infected regions in the spatial transcriptomics data (Figure 3H). The infected regions had a much higher percentage of ASR reads compared to the surrounding regions (Figure 3H), suggesting that the surrounding regions had a low presence of ASR. Although the surrounding regions had a significantly higher percentage of ASR reads compared to the healthy regions, the average percentage of ASR reads in the surrounding regions was very close to that in the healthy regions while drastically lower compared to the infected regions (Figure 3H). These results suggest that the surrounding regions indeed had a very low presence of the ASR pathogen. Next, we examined the expression of defense response-related genes in the surrounding and infected regions (Figure 3I). We combined the genes from the GO term “defense response” as a gene module and plotted the expression of these genes as a module score in the healthy, surrounding, and infected regions (Figure 3I, Table S2). We found that although both surrounding and infected regions had higher expression of this defense response gene module compared to the healthy regions, the surrounding regions had significantly higher expression of the defense response genes than the infected regions (Figure 3I, Table S2). These results suggest that although the surrounding regions had a much lower presence of ASR than the infected regions, the surrounding regions had higher expression of defense response genes.

To further verify the defense-related gene expression pattern in the surrounding and infected regions, we examined the expression of known ASR-induced genes from the literature and found that the ASR-induced genes had higher expression in the surrounding regions compared to the infected regions (Figure S3E, Table S2) (Morales et al., 2013; Cabre et al., 2021), further confirming that the surrounding regions had higher activation of defense-related genes than the infected regions. Moreover, when we plotted the expression of the ASR-induced genes across spots, we observed clear enrichment of these genes in the surrounding regions and less enrichment in the infected regions (Figure S3F, Table S2). Consistently, the expression of the ASR-induced genes showed enriched spots on the sections (Figure S3G, Table S2), and these enriched spots overlapped with the surrounding regions instead of the infected regions, which is evident in the paradermal sections (Figure 2E and S3G, Table S2). This further supports that the surrounding regions bordering the infected regions had higher expression of defense response-related genes than the infected regions.

To better understand the differences between the infected regions and the surrounding regions, we performed DEG analysis by directly comparing the infected regions to the surrounding regions (Figure S3H). Using the DEGs derived from this comparison, we further performed GO term analysis and found that many defense response-related GO terms, such as “defense response to other organism” and “flavonoid biosynthetic process,” were enriched in the downregulated genes in the infected regions compared to the surrounding regions (Figure 3J, Table S1). This suggests that the defense response in the infected regions is indeed suppressed compared to the surrounding regions. Altogether, we revealed that the surrounding regions and the infected regions correspond to two distinct cell states during the plant response to pathogen infection, with the surrounding regions bordering the infected regions. The surrounding regions had a much lower presence of the pathogen than the infected regions but had higher expression of defense response-related genes. These results further underscore the importance of using spatial transcriptomic techniques to decipher the spatial organization of the plant immune response to pathogen infection.

### 3.6 Gene co-expression network analysis identifies key immune responsive gene module in stressed cells

The plant defense response to pathogens involves complex changes at the whole transcriptome level, encompassing many different transcriptional pathways and potentially various waves of gene expression activation and suppression (Dobon et al., 2016). To delineate the plant defense response at the systems biology level using single-cell transcriptomics data, we constructed a single-cell gene co-expression network through hdWGCNA using the mesophyll cells in our snRNA-seq data (Figure 4A and S4A) (Morabito et al., 2023). This co-expression network was constructed using both the healthy and stressed mesophyll cells annotated in the snRNA-seq dataset (Figure 4A and S4A), facilitating the identification of gene co-expression modules in the stressed mesophyll cells. The mesophyll co-expression network analysis identified nine different co-expression gene modules, labeled from M1 to M9 (Figure 4A and S4A). UMAP visualization of all gene modules revealed the relative relationship between individual modules and showed that module M3 and module M4 were the two largest gene modules in the network (Figure 4A).

**Figure 4.**
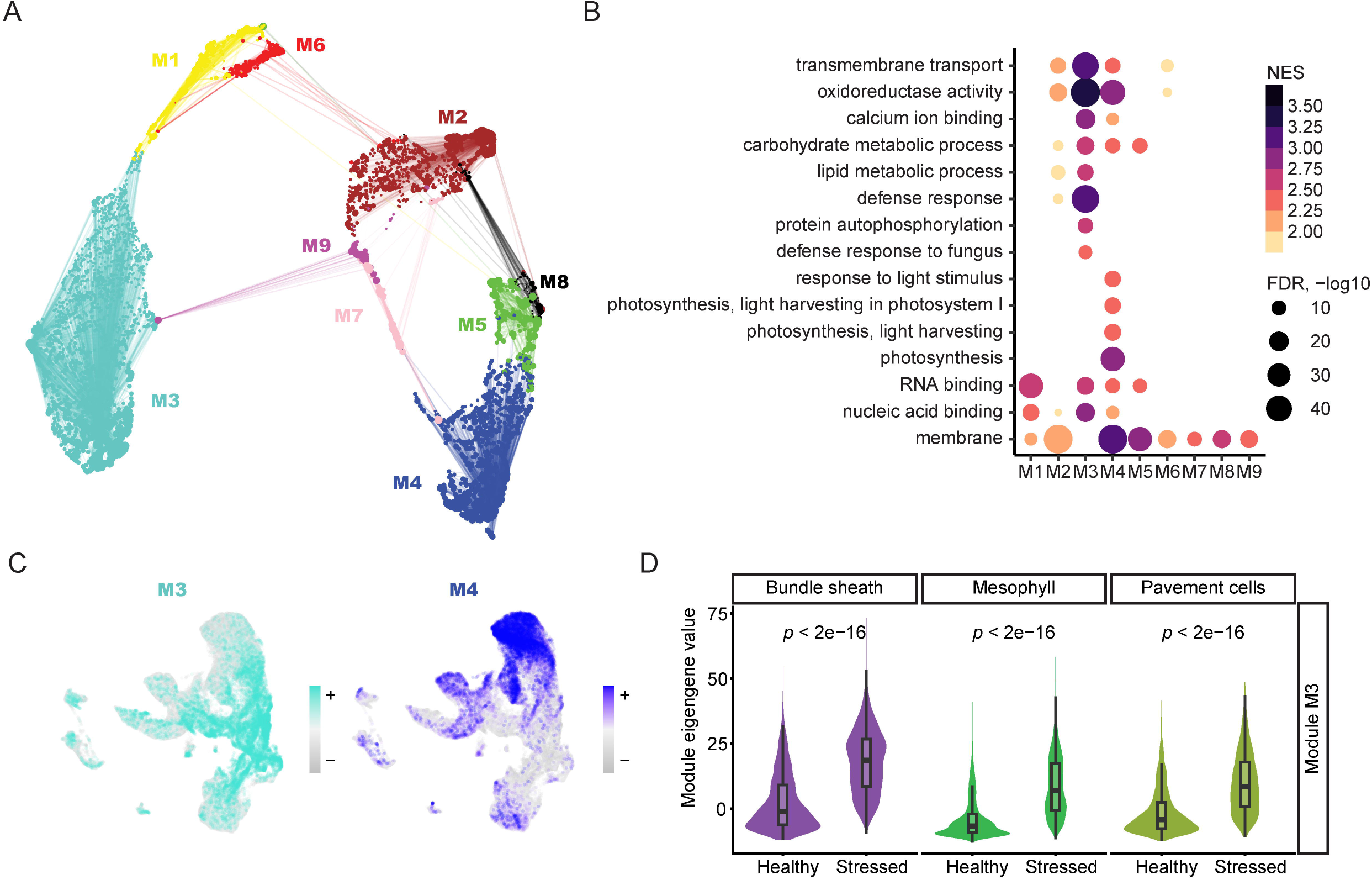
Gene co-expression network analysis with single cell resolution captures key defense response-related gene module. **A)** Gene-centric UMAP of mesophyll co-expression modules constructed with hdWGCNA using snRNA-seq data. Dots indicate genes, and lines indicate strength of relationship with module hub genes. **B)** Dot plot of enriched GO terms associated with genes by mesophyll co-expression module membership. Size of dots corresponds to −log10 of adjusted *p* value, colored by NES. **C)** Module eigengene values for modules M3 and M4 plotted in snRNA-seq UMAP. **D)** Comparison of M3 module eigengene distributions in stressed and healthy cells across three cell types in snRNA-seq. Significance was determined via an unpaired two-sided Wilcoxon Rank Sum test.

To further understand the biological significance of the different gene modules, we performed GO term analysis using the genes assigned to each module (Figure 4B, Table S1). The GO term analysis revealed that module M3 included many genes involved in defense response-related GO terms, such as “defense response to fungus,” while module M4 encompassed many genes involved in photosynthesis-related GO terms, such as “photosynthesis, light harvesting” (Figure 4B, Table S1). These results suggest that module M3 is an immune response-related module, while module M4 is a photosynthesis-related module. To further understand the expression of the various modules identified in the co-expression network, we plotted the expression of the eigengenes of each module across all nuclei (Figure 4C and S4B). Interestingly, module M3 had enriched expression in the stressed cells in the snRNA-seq dataset (Figure 4C), indicating that module M3 is activated in the stressed cells, in line with the observation that defense response-related GO terms are enriched in the genes assigned to module M3. Moreover, module M4 had enriched expression in the healthy mesophyll cells in the snRNA-seq dataset (Figure 4C), agreeing with the observation that photosynthesis-related GO terms were enriched in the genes assigned to module M4.

Next, we performed differential module eigengene analysis, examining the eigengenes for modules M3 and M4 and comparing their expression in stressed cells versus healthy cells (Figure 4D and S4C). We found that the eigengene of module M3 was upregulated in the stressed mesophyll cells compared to healthy mesophyll cells (Figure 4D), indicating that module M3 is indeed an immune response-related gene module. Interestingly, even though module M3 is derived from mesophyll cells, its eigengene was upregulated in stressed bundle sheath and pavement cells compared to their healthy counterparts (Figure 4D), suggesting that different cell types share similar transcriptional changes in response to pathogen infection. Additionally, we performed differential module eigengene analysis for the photosynthesis-related module M4 (Figure S4C). We found that the eigengene of module M4 was downregulated in stressed mesophyll cells compared to healthy mesophyll cells (Figure S4C), verifying the finding that photosynthesis-related pathways are downregulated upon pathogen infection (Bilgin et al., 2010). Altogether, we identified a key immune response gene module that is highly induced in the stressed cells in the snRNA-seq data, deepening our understanding of the sophisticated transcriptional network involved in the plant defense response to pathogen invasion.

## 4 Discussion

In this study, using snRNA-seq and spatial transcriptomics, we have characterized the spatially organized soybean plant response to ASR pathogen infection. The spatial and single-nuclei transcriptomics technologies used in this study allowed the simultaneous capture of the host and pathogen transcriptomes, enabling the identification of infection sites within the soybean leaf sections. Furthermore, the high resolution of the single-nuclei transcriptomics and the annotation of snRNA-seq clusters facilitated the annotation of clusters in the spatial transcriptomics data. The functional annotation of the spatial transcriptomics data led to the important finding that the host immune response is spatially organized into two different cell states/regions: the infected regions at the center of pathogen infection exhibiting a relatively weak defense response, and the surrounding regions bordering the infected regions mounting a stronger defense response with minimal pathogen contact. Finally, through gene co-expression network analysis, we identified a defense response-related gene module that is induced in the stressed cells identified in the snRNA-seq. Together, the findings in this study provide new evidence supporting the recently emerged hypothesis that localized acquired resistance in the surrounding regions of the infection site could be important for pathogen containment, despite the host cells in the infection center being immune-suppressed by effectors secreted from the pathogen.

### 4.1 Localized acquired resistance

The concept of localized acquired resistance was first proposed in 1961 by A. F. Ross, with the observation that a 1-2 mm zone surrounding the tobacco mosaic virus infection lesion developed resistance to subsequent inoculation of tobacco mosaic virus in the host species *Nicotiana tabacum*. However, the mechanism and importance of this localized acquired resistance remained elusive (Ross, 1961). Recently, a study revisited the concept of localized acquired resistance and proposed that its induction is a key component of the mechanism through which effector-triggered immunity circumvents effector-triggered susceptibility (Jacob et al., 2023a). This newly proposed spatially organized mechanism of plant immune response is supported by another recently published study where single-cell and spatial transcriptomics technologies were applied to study *Arabidopsis* immune response to bacterial infection (Nobori et al., 2025). In their study, two cell populations were identified in plant immunity: the primary immune responder (PRIMER) cells and the bystander cells. The PRIMER cells had the strongest association with the pathogen and were surrounded by the bystander cells. Similarly, in our study, we identified two distinct cell states/regions that are spatially organized at or near the ASR infection site. On the one hand, the infected regions correspond to the PRIMER cells, with the presence of the ASR pathogen, and have a weaker defense response than the surrounding regions, in line with the proposed pathogen effector-triggered suppression of immune response (Jacob et al., 2023a). On the other hand, the surrounding regions resemble the bystander cells with localized acquired resistance. We observed that the surrounding regions had minimal presence of *P. pachyrizi* but a stronger defense response compared to the infected regions. This observation aligns with the hypothesis by Jacob et al. that the infected cells release an uncharacterized signaling molecule that travels to the surrounding regions to activate a cell non-autonomous defense response (Jacob et al., 2023a). Therefore, our study provides important further evidence that the plant immune response to pathogen invasion involves two spatially organized regions: the infected regions in contact with the pathogen and the surrounding regions bordering the infected regions that exhibit cell non-autonomous localized acquired resistance.

### 4.2 Spatial and single cell transcriptomics of both host and pathogen transcriptome

Plant-microbe interaction is an important field of study in crops, encompassing interactions between plants and beneficial microbes to form root nodules, as well as interactions between plants and various disease-triggering pathogens (Zhu et al., 2023b; Hu et al., 2024; Serrano et al., 2024b). During these plant-microbe interactions, the plant response is usually very heterogeneous (Zhu et al., 2023b; Hu et al., 2024; Serrano et al., 2024b). Depending on whether the plant cells are directly contacting the microbes, their responses can vary significantly (Betsuyaku et al., 2018). Moreover, during plant-pathogen interaction, the plant cells in contact with the pathogen and the plant cells surrounding the infection site can mount very different spatially organized responses (Jacob et al., 2023b). Therefore, cell-specific high resolution and spatial information are crucial aspects of characterizing plant-microbe interactions (Zhu et al., 2023b; Hu et al., 2024; Serrano et al., 2024b).

Many single-cell/nuclei RNA-seq studies have been conducted to study plant-microbe interactions (Cao et al., 2023; Delannoy et al., 2023; Liu et al., 2023b; Tang et al., 2023; Cervantes-Perez et al., 2024; Pereira et al., 2024; Yan et al., 2025), but spatial transcriptomics studies are less prevalent (Liu et al., 2023a; Serrano et al., 2024a; Nobori et al., 2025). Similar to another study that used spatial transcriptomics technology to capture host response to pathogen infection (Nobori et al., 2025), our current study captured the host and pathogen transcriptomes in the same spatial transcriptomics experiment. This technical aspect of our study is especially important as it enabled us to identify specific regions of pathogen infection within the tissue sections sampled for the spatial transcriptomics experiment. It also allowed us to delineate the surrounding and infected regions, which play different roles in the plant immune response.

Our snRNA-seq data also captured both the host and pathogen nuclei. However, the total number of pathogen nuclei captured was fewer than 100, which is not enough to perform conclusive statistical analysis. This low number of captured pathogen nuclei could be related to the relatively early stages of infection, up to 5 DPI. Future studies could expand the time course to later infection stages to capture more pathogen nuclei for transcriptome examinations. Nevertheless, the transcriptomic response of the pathogen in the early stages of infection could be of interest as well (Gervais et al., 2017). Therefore, future studies could focus on using enrichment techniques, such as the recently published Programmable Enrichment via RNA FlowFISH by sequencing (PERFF-seq) (Abay et al., 2025), to enrich for pathogen nuclei and study their transcriptomic activities.

In our snRNA-seq data, we observed that the percentage of stressed cells was highest at 1 DPI, slightly dropped at 3 DPI, and then increased again at 5 DPI (Figure S2E). This observation aligns with the reported biphasic transcriptional response to ASR infection in soybean leaves (van de Mortel et al., 2007; Schneider et al., 2011; Chicowski et al., 2024). The 1 DPI time point processed in our study could correspond to the initial phase of transcriptional changes, while the 3 DPI time point resembles the middle quiescent period of the transcriptional response, and the 5 DPI time point indicates a second phase of transcriptional response. Future in-depth examination of this biphasic transcriptional response with single-cell resolution could deepen our understanding of the temporal and spatial plant defense response to pathogen infection.

### 4.3 Gene co-expression networks for immune response at single cell resolution

Plant responses to pathogen infection encompass complex regulatory pathways, including PAMP-triggered immunity (PTI) activated by cell surface-localized pattern recognition receptors and effector-triggered immunity (ETI) initiated by intracellular nucleotide-binding leucine-rich repeat receptors (Ngou et al., 2022). PTI and ETI pathways function synergistically to activate similar pathways, such as calcium signaling, reactive oxygen species (ROS) burst, mitogen-activated protein kinase (MAPK) signaling pathway, and transcriptomic activation of defense-related genes (Dodds et al., 2024). The transcriptomic response itself is sophisticated and may constitute multiple waves of transcriptional regulation, such as the transcriptional activation of a transcription factor, which in turn activates or represses the expression of other defense response-related genes (Dobon et al., 2016).

The advent of transcriptomics technologies has enabled the simultaneous examination of changes at the whole transcriptome level in plant responses to pathogen infection (van de Mortel et al., 2007; Wise et al., 2007), but the analysis is often performed by comparing different conditions and focusing on genes with the most extreme expression changes (Peyraud et al., 2017). This approach is sometimes not fruitful due to the redundancy of the biological gene network, with redundancy of various genes and different signaling pathways. Because of these redundancies, modifying the function of individual genes may not be sufficient to generate the desired disease resistance phenotype at the whole organism level (Peyraud et al., 2017). Therefore, modeling the gene network for plant immunity and identifying gene modules important for the immune response can provide great insights into the sophisticated gene network in host cells that are responsive to pathogen infection.

So far, studies of plant pathogen interactions have built gene networks using bulk transcriptome data or at the cell type-specific level (Yuan et al., 2018; Yue et al., 2024), but plant immunity gene networks with high cell resolution remain limited (Cao et al., 2023). In our current study, we have built a gene co-expression network for soybean cells in response to ASR pathogen infection using an approach that provides high cell resolution to reveal more granular gene co-expression relationships. Importantly, combined with our carefully annotated snRNA-seq data, we showed that the immune response-related gene co-expression module we identified is indeed highly enriched and induced in the stressed soybean cell population. This identification of this specific immune response module serves as a foundation for future studies examining gene networks for plant immune responses and opens the door for examination of the effects of gene perturbations on disease resistance at the gene network level.

## Acknowledgements

We thank Bailin Li for his scientific suggestions and guidance on this manuscript. We also thank Jeff Habben for his support with this work. We thank Katherine Thilges and Heather Cartwright for their help with histology and microscopy. We thank Charlotte Harris and John Tenhundfeld for sequencing.

