## Supplementary Figure 1 for "Spatial and single-cell transcriptomics capture two distinct cell states in plant immunity"

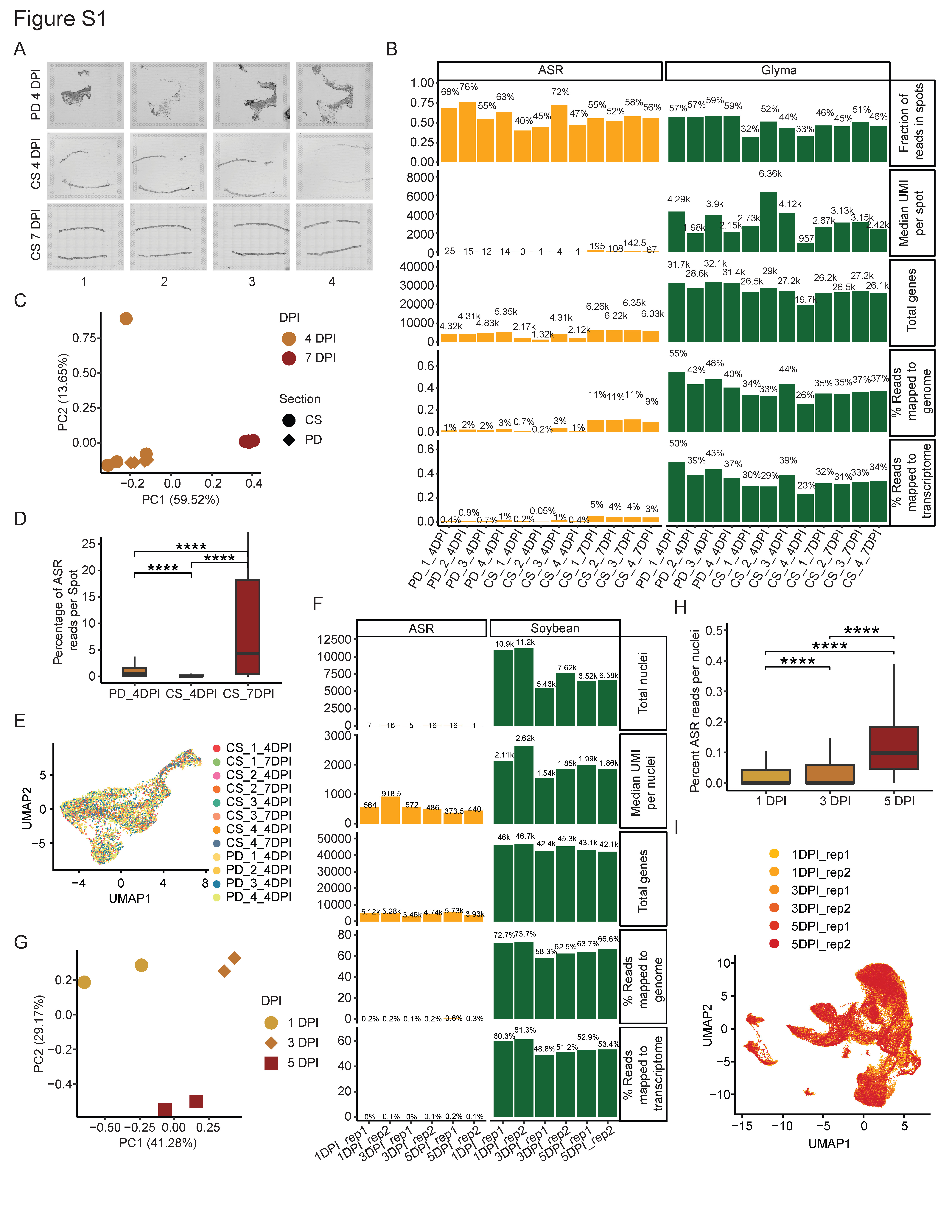
**Supplemental Figure 1. A)** Brightfield uncropped images of leaf sections used for spatial analysis. **B)** Sequencing metrics for spatial analysis separated by genome. **C)** PCA of pseudo-bulked normalized counts for spatial analysis. **D)** Percent ASR reads per spot measured across all spatial transcriptomics samples. * indicates *p* < 0.05, ** indicates *p* > 0.01, *** indicates *p* < 0.001, **** indicates *p* < 0.0001. **E)** UMAP of all spots from all sections. **F)** Sequencing metrics for snRNA-seq separated by genome. **G)** PCA of pseudo-bulked counts for snRNA-seq. **H)** Percent ASR reads per nuclei measured across days of ASR infection. * indicates *p* < 0.05, ** indicates *p* > 0.01, *** indicates *p* < 0.001, **** indicates *p* < 0.0001. **I)** UMAP of all nuclei from all snRNA-seq samples.
