## Supplementary Figure 2 for "Spatial and single-cell transcriptomics capture two distinct cell states in plant immunity"

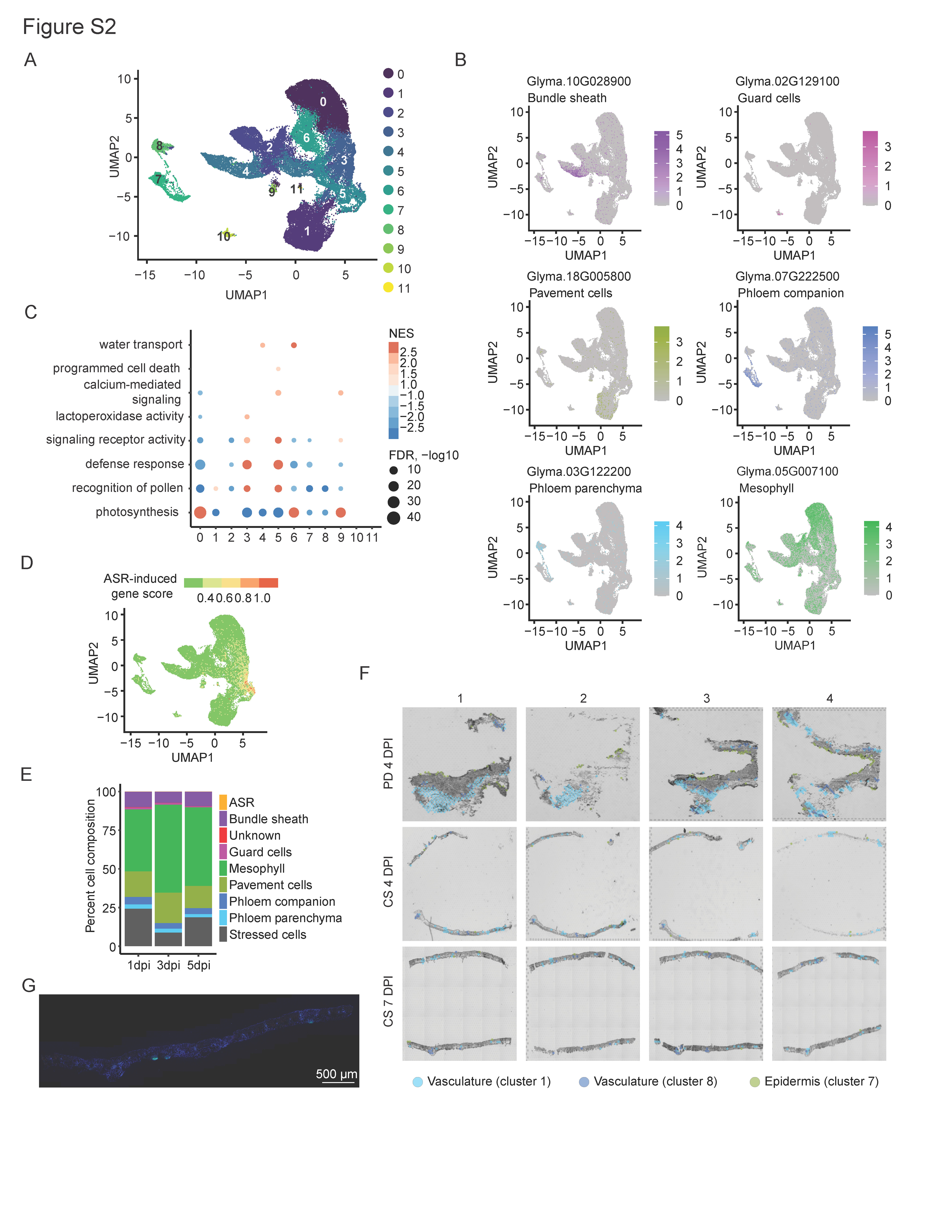
**Supplemental Figure 2. A)** UMAP of snRNA clusters. **B)** Expression of known leaf cell type marker genes in snRNA-seq dataset. **C)** Dot plot of enriched GO terms associated with each snRNA-seq cluster’s marker genes. Size of dots corresponds to -log10 of adjusted *p* value, colored by Normalized Enrichment Score (NES). **D)** ASR-induced gene expression described as a single score across snRNA UMAP. **E)** Cell type composition in snRNA across infection time. **F)** Spatial clusters 1 and 8 (blue) overlapping with midrib vascular bundles in cross section as well as visible veins in paradermal sections. Spatial cluster 7 (green) outlines paradermal sections and few external spots in cross sections. G) Fluorescence image of 7 DPI cross section. Green fluorescence shows localization of fungal cells stained with wheat germ agglutinin Alexa Fluor 488 conjugate. Scale bar is 500 μm.
