## Supplementary Figure 3 for "Spatial and single-cell transcriptomics capture two distinct cell states in plant immunity"

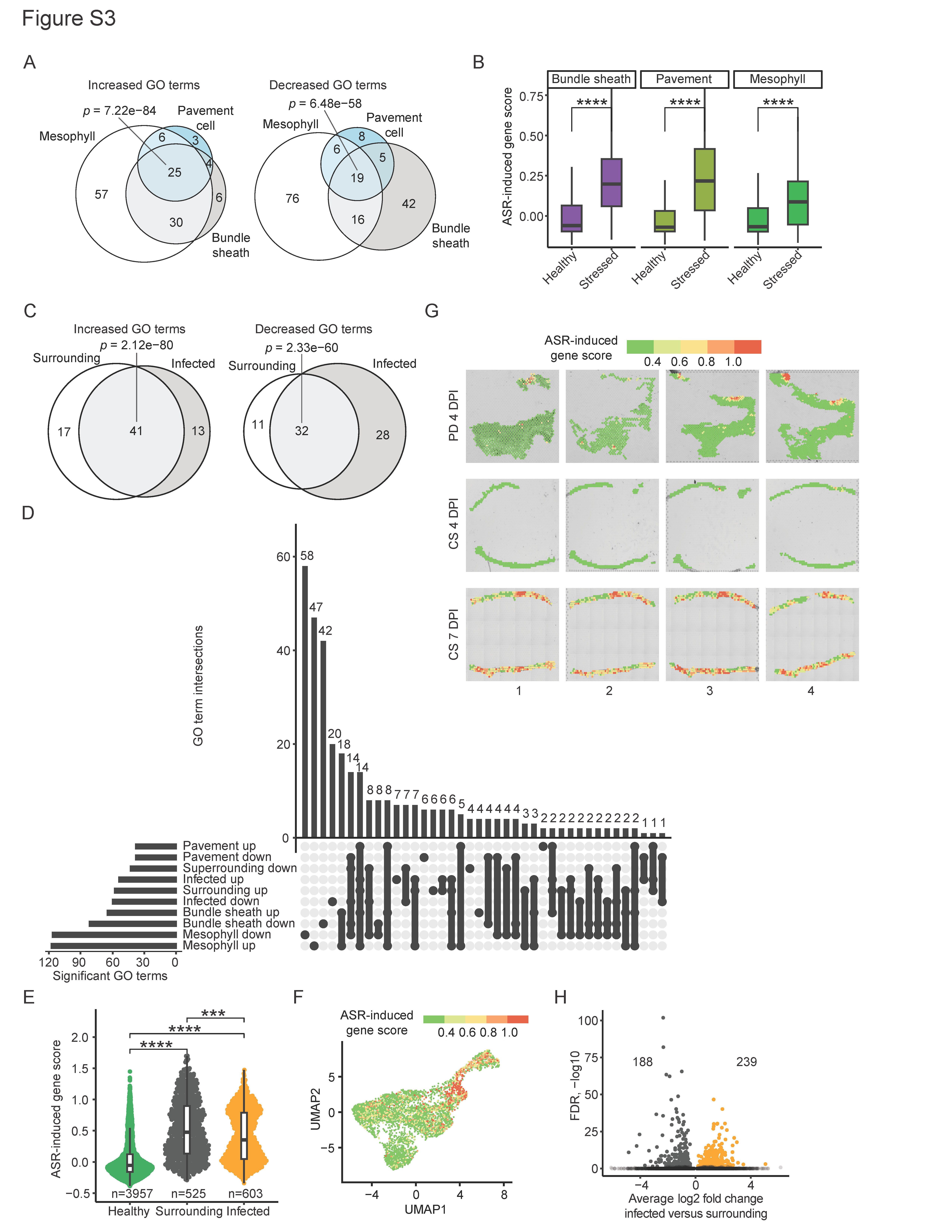

**Supplemental Figure 3. A)** Venn diagrams showing agreement in significantly enriched and depleted GO terms in stressed cell types. Significance determined by super exact test. **B)** ASR-induced gene expression described as a single score per nuclei in each cell type cluster or stressed subcluster. * indicates *p* < 0.05, ** indicates *p* > 0.01, *** indicates *p* < 0.001, **** indicates *p* < 0.0001. **C)** Venn diagrams showing agreement in significantly enriched and depleted GO terms in surrounding and infected spatial regions compared to healthy regions, respectively. Significance determined by hypergeometric test. **D)** Upset plot comparing enriched GO terms for stressed versus healthy nuclei in each snRNA-seq cell type and for surrounding and infected spatial regions. **E)** ASR-induced gene expression described as a single score per spot in each spatial region. * indicates *p* < 0.05, ** indicates *p* > 0.01, *** indicates *p* < 0.001, **** indicates *p* < 0.0001. **F)** ASR-induced gene expression described as a single score across all spatial spots. **G)** ASR-induced gene expression described as a single score across all spots on all sections. **H)** Volcano plots of significantly differentially expressed soybean genes (padj < 0.05) in infected vs surrounding spatial regions.
