## Supplementary Figure 4 for "Spatial and single-cell transcriptomics capture two distinct cell states in plant immunity"

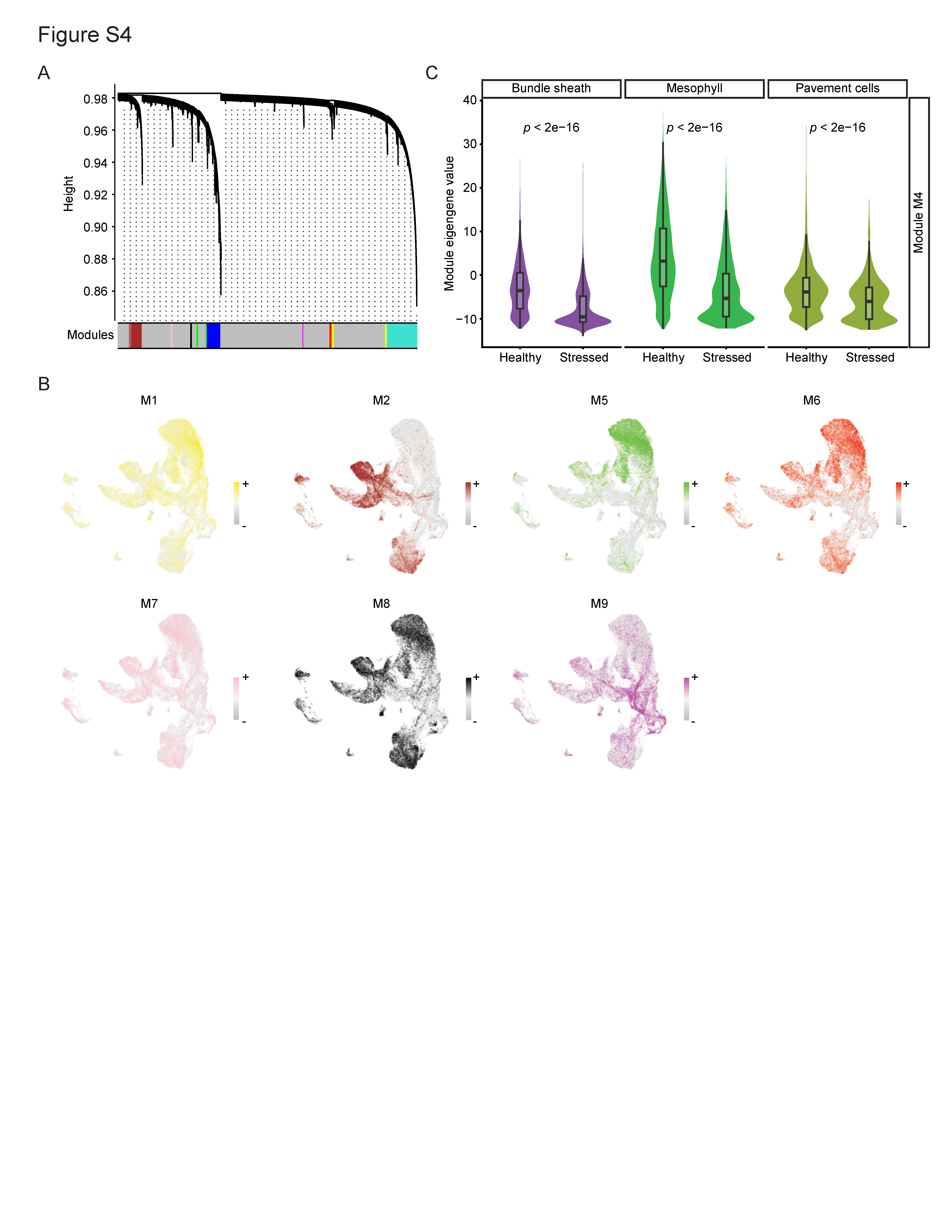
**Supplemental Figure 4. A)** Dendrogram of co-expression network modules constructed by hdWGCNA on mesophyll cells using snRNA-seq data. Colored bar indicates module assignment for each gene. **B)** Module eigengene values for M1, M2, and M5-9 plotted in snRNA-seq UMAP. **C)** Comparison of M4 module eigengene distributions in stressed and healthy cells across three cell types in snRNA-seq. Significance was determined via an unpaired two-sided Wilcoxon Rank Sum test.
